# A pseudorabies virus serine/threonine kinase, US3, promotes retrograde transport in axons via Akt/mToRC1

**DOI:** 10.1101/2021.10.12.464169

**Authors:** Andrew D. Esteves, Orkide O. Koyuncu, Lynn W. Enquist

**Author notes:** **Correspondence:** Andrew D. Esteves, Lynn W. Enquist. Department of Microbiology and Molecular Genetics, School of Medicine, University of California, Irvine, Irvine, CA, USA.

## Abstract

Infection of peripheral axons by alpha herpesviruses (AHVs) is a critical stage in establishing a life-long infection in the host. Upon entering the cytoplasm of axons, AHV nucleocapsids and associated inner-tegument proteins must engage the cellular retrograde transport machinery to promote the long-distance movement of virion components to the nucleus. The current model outlining this process is incomplete and further investigation is required to discover all viral and cellular determinants involved as well as the temporality of the events. Using a modified tri-chamber system, we have discovered a novel role of the pseudorabies virus (PRV) serine/threonine kinase, US3, in promoting efficient retrograde transport of nucleocapsids. We discovered that transporting nucleocapsids move at similar velocities both in the presence and absence of a functional US3 kinase; however fewer nucleocapsids are moving when US3 is absent and move for shorter periods of time before stopping, suggesting US3 is required for efficient nucleocapsid engagement with the retrograde transport machinery. This led to fewer nucleocapsids reaching the cell bodies to produce a productive infection 12hr later. Furthermore, US3 was responsible for the induction of local translation in axons as early as 1hpi through the stimulation of a PI3K/Akt-mToRC1. These data describe a novel role for US3 in the induction of local translation in axons during AHV infection, a critical step in transport of nucleocapsids to the cell body.

**Importance:** Neurons are highly polarized cells with axons that can reach centimeters in length. Communication between axons at the periphery and the distant cell body is a relatively slow process involving the active transport of chemical messengers. There’s a need for axons to respond rapidly to extracellular stimuli. Translation of repressed mRNAs present within the axon occurs to enable rapid, localized responses independently of the cell body. AHVs have evolved a way to hijack local translation in the axons to promote their transport to the nucleus. We have determined the cellular mechanism and viral components involved in the induction of axonal translation. The US3 serine/threonine kinase of PRV activates Akt-mToRC1 signaling pathways early during infection to promote axonal translation. When US3 is not present, the number of moving nucleocapsids and their processivity are reduced, suggesting that US3 activity is required for efficient engagement of nucleocapsids with the retrograde transport machinery.

## Introduction

Members of the alpha herpesvirus (AHV) subfamily, including the human pathogens, herpes simplex virus type-1 and 2 (HSV-1 and HSV-2), as well as the animal pathogen, pseudorabies virus (PRV), are pantropic viruses capable of infecting the peripheral (PNS) and central (CNS) nervous system of their hosts. Infection of the highly polarized PNS at axon terminals occurs after infection of the epithelial layer. Once in the axonal cytoplasm, nucleocapsids undergo efficient retrograde transport to the nucleus where the viral DNA is transcribed and either a productive or a life-long latent infection is established. In natural hosts, AHVs tend to establish a latent infection. Reactivation from latency results in the anterograde transport of progeny virion particles in the axon to re-infect the epithelial layer that promotes spread to new hosts. In rare events, progeny virion particles can spread in the opposite direction and transsynaptically invade the CNS, often leading to death of the organism. In non-natural hosts, a productive infection followed by invasion of the CNS and death are the most common ^1^.

The recruitment of the retrograde transport machinery to mediate the active transport of virion particles is facilitated by the viral nucleocapsid and inner-tegument proteins (UL36, UL37, and US3) ^2,3^. This process, while dispensable in non-polarized cells, is essential for neuronal infection via axons due to the large distance between the axon terminal and the cell body ^3–5^. Despite this, the viral and cellular factors involved, and the temporality of the events are not well understood. Following fusion of the viral and cellular membranes, a breach in the actin cytoskeleton is created through the activation of cofilin by the US3 protein ^6^. The microtubule plus-tip proteins, EB-1 and CLIP170 also have been shown to aid in this process during HSV-1 infection ^7^. The UL36 inner-tegument protein then interacts with dynactin, to recruit the nucleocapsid complex to dynein, the retrograde-directed motor protein ^8,9^. UL37, even though it has not been shown to interact with dynein, is capable of modulating its activity ^4^. We have previously shown that upon infection of axons with PRV, translation of a subset of axonally-localized mRNA occurs, producing a subset of proteins related to intracellular transport and cytoskeletal remodeling ^10^. Among these was the dynein regulator, Lis1. Local translation in axons was shown to be essential for the efficient transport of virion components through the axon; however the mechanism that regulates this is unknown.

Translation of cellular mRNA is a tightly regulated process, with the initiation stage being the most rate-limiting. The PI3K/Akt-mToRC1 signaling pathway is the canonical route by which translation initiation occurs in eukaryotic cells ^11^. AHVs have been demonstrated to manipulate this signaling pathway to support infection and replication. HSV-2 infection induces Akt phosphorylation upon binding of gB, on the virion envelope, to α_v_β_3_ integrins leading to release of intracellular calcium stores to promote entry of nucleocapsids into the cell ^12,13^. In HSV-1 infected cells activation of mToRC1 was induced by the phosphorylation of Akt substrates in an Akt independent manner. This work demonstrated that a viral kinase could act as an Akt surrogate to bypass cellular signal pathway control mechanisms to promote constitutive viral replication ^14^. In VZV infected cells Akt phosphorylation is increased and required for efficient replication^15–17^. PRV infection induces Akt phosphorylation to mediate anti-apoptosis effects on infected cells ^18^. Despite these findings in non-neuronal cells, it is not known if AHV infection of axons induces Akt signaling pathways. In this paper we investigated whether PRV infection stimulates the Akt-mToRC1 signaling pathway to induce local translation in axons.

US3 is a multifunctional, viral encoded serine/threonine kinase present in the inner-tegument layer of the virion and one of only two protein kinases conserved by all AHVs ^19^. Although US3 has been shown to be dispensable for virus replication in cell culture, it is vital for viral fitness *in vivo* ^20–25^. Some of the most notable functions of US3 include the inhibition of apoptosis in infected cells through the activation of Akt and NF-κB signaling pathways, promotion of nuclear egress of newly-made nucleocapsids, and disassembly of actin stress fibers by cofilin activation ^26–28^. Due to its serine/threonine kinase function, known interactions with Akt, and its presence in the tegument layer, and thus delivered directly to the cytoplasm, we hypothesized that US3 stimulates Akt-mToRC1 signaling pathways in axons early after infection, to induce local translation.

In this study, we established a novel role for PRV US3 in the induction of translation in axons via a PI3K/Akt-mToRC1 signaling pathway early after virion entry to promote the efficient retrograde transport of nucleocapsids to the cell body. In the absence of US3 or Akt phosphorylation, the number of transporting nucleocapsids and their processivity were significantly reduced. These events led to a reduction in the number of infected neuronal cell bodies later in infection. Together, these findings suggest a role for US3 in the continuous engagement of PRV nucleocapsids with the retrograde transport machinery. Due to the significance this stage of infection plays in AHV infection of neurons at the axon, US3 may serve as a target for drugs aiming to prevent the life-long, reactivatable infection caused by AHVs.

## Results

### Akt is phosphorylated in axons early after PRV infection

To determine if Akt is phosphorylated in axons early after PRV infection, we cultured primary rat superior cervical ganglion (SCG) neurons *in vitro* in Campenot tri-chambers (Fig.1A). This cell culture system allows for the fluidic separation of neuronal cell bodies (in the S compartment) from axons (in the N compartment), enabling us to infect pure populations of axons and monitor responses independently of the cell bodies ^29^. N compartment axons were infected with 10^6^ plaque forming units (PFU) of a PRV-Becker recombinant expressing a monomeric red fluorescent protein (mRFP)-tagged capsid protein (VP26) termed PRV 180. Akt phosphorylation was monitored using a phospho (p-) S^473^-Akt antibody. Phosphorylation at serine 473 is the most well-known mechanism of Akt activation ^30^ and can be observed as early as 30 minutes post infection (mpi) in axons and continues to at least 180mpi (Fig.1B), indicating that PRV infection does induce Akt phosphorylation in axons. Furthermore, when axons in the N compartment were pretreated with LY294,002 (a potent and selective PI3K inhibitor) prior to infection, no Akt phosphorylation was observed, suggesting that PRV-induced Akt phosphorylation is PI3K-dependent (Fig.1C). When axons were pretreated with the mToRC1 inhibitor rapamycin, or the translation inhibitor cycloheximide, Akt phosphorylation did occur (Fig.1C). No change in Akt phosphorylation occurred in S compartment cell bodies 1hour post infection (hpi) when the N compartment was infected, demonstrating that the infection and intracellular signals do not reach the cell body in that time (Fig.1D).

**Fig 1.**
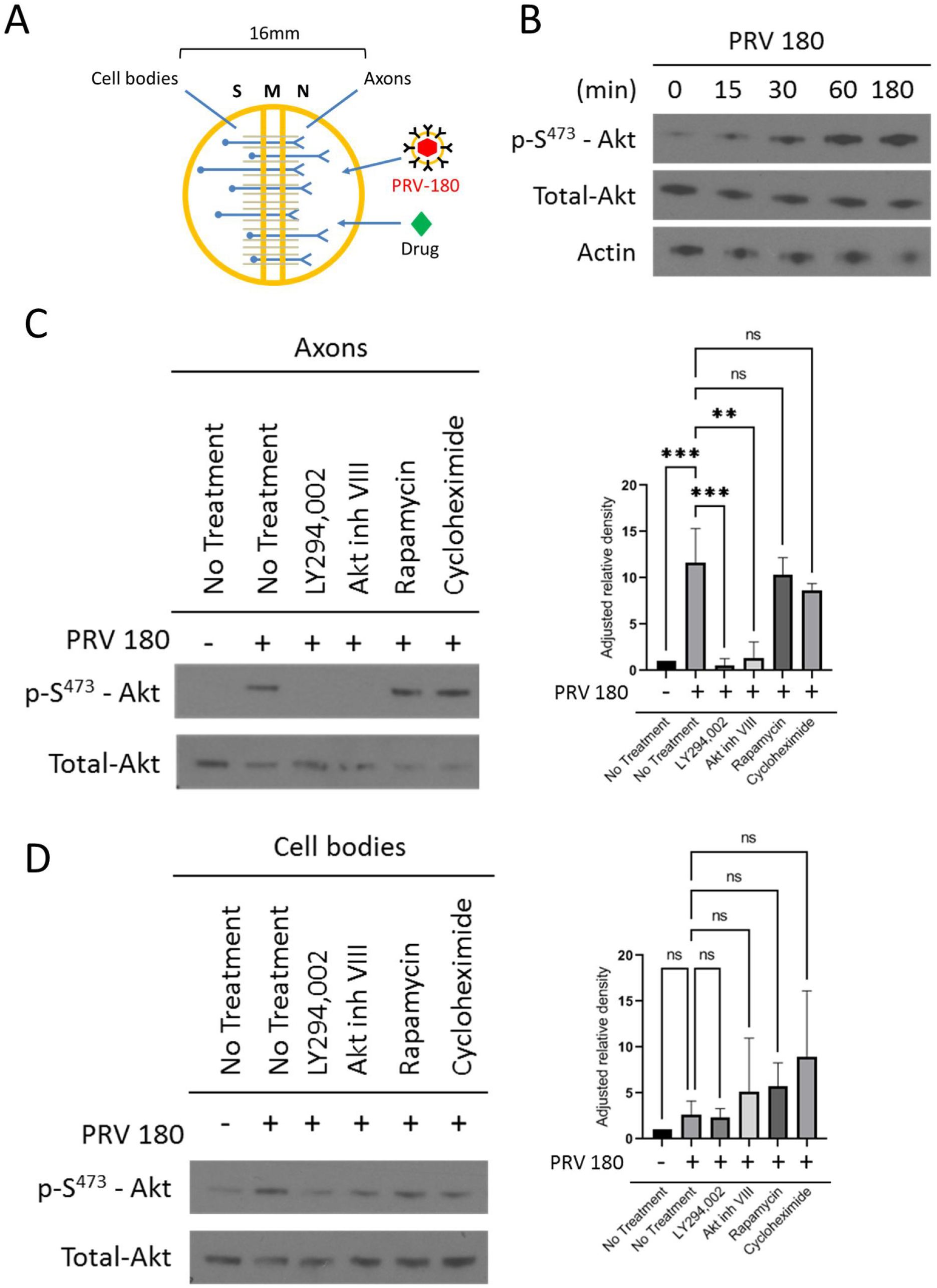
Akt is phosphorylated in axons during PRV infection. (A) A Campenot tri-chamber neuronal culture system divided into soma (S), middle (M), and neurite (N) compartments used to separate neuronal cell bodies from axons. Addition of virus or drug into the N compartment allows for the study of axonal responses independent of the cell body. (B and C) Immunoblot of p-S^473^-Akt in axons infected with PRV 180 in the N compartment. (B) Infections continued for 0min, 15min, 30min, 60min, and 180min. (C and D) N compartments were pretreated with LY294,002, Akt inhibitor VIII, rapamycin, or cycloheximide prior to infection. (C) N compartments were harvested 1hpi. (D) S compartments were harvested 1hpi in the N compartment. (C and D) p-S^473^-Akt bands were normalized to total-Akt bands using a densitometry assay. Mean ± Std Dev with n = 3 for each condition are plotted with ** p ≤ 0.01, *** p ≤ 0.001, using a one-way ANOVA (ns = not significant).

### Akt phosphorylation in axons is required for efficient retrograde transport of PRV nucleocapsids

Next, we determined whether infection-induced Akt phosphorylation in axons affected the spread of PRV to the distant cell body. N compartments were treated with a green lipophilic dye, Fast-DiO, to label the membranes of all axons in the N compartment and their attached cell bodies in the S compartment such that only the cell bodies with axons extended into the N compartment were labeled. 12 hours post DiO staining, N compartments were infected with PRV 180 (Fig.2A). At 12hpi, infected cell bodies were visualized by the presence of red fluorescence from the mRFP-VP26 fusion protein (Fig.2B). The presence of dual-colored cell bodies (expressing red and green fluorescence) were those who were directly infected via axons in the N compartment. The ratio of dual-colored cell bodies to total green cell bodies represents the efficiency of retrograde infection. When N compartment axons were pretreated with Akt inhibitor VIII for 1 hour prior to infection, we observed a ∼56.4% ± 14.7% reduction in the number of dual-colored cell bodies compared to PRV 180 infection alone. Additionally, when axons were pretreated with LY294,002, rapamycin, or cycloheximide, dual-colored cell bodies were reduced by ∼48% ± 16.2%, ∼58% ± 17.6%, and ∼65.7 ± 9.5% respectively. Ras/MAPK is also known to promote translation by signaling through mToRC1^31–36^, however pretreatment of axons with the Erk1/2 inhibitor, U0126, had no significant effect on retrograde infection (∼8.3% ± 14%) (Fig.2C).

**Fig 2.**
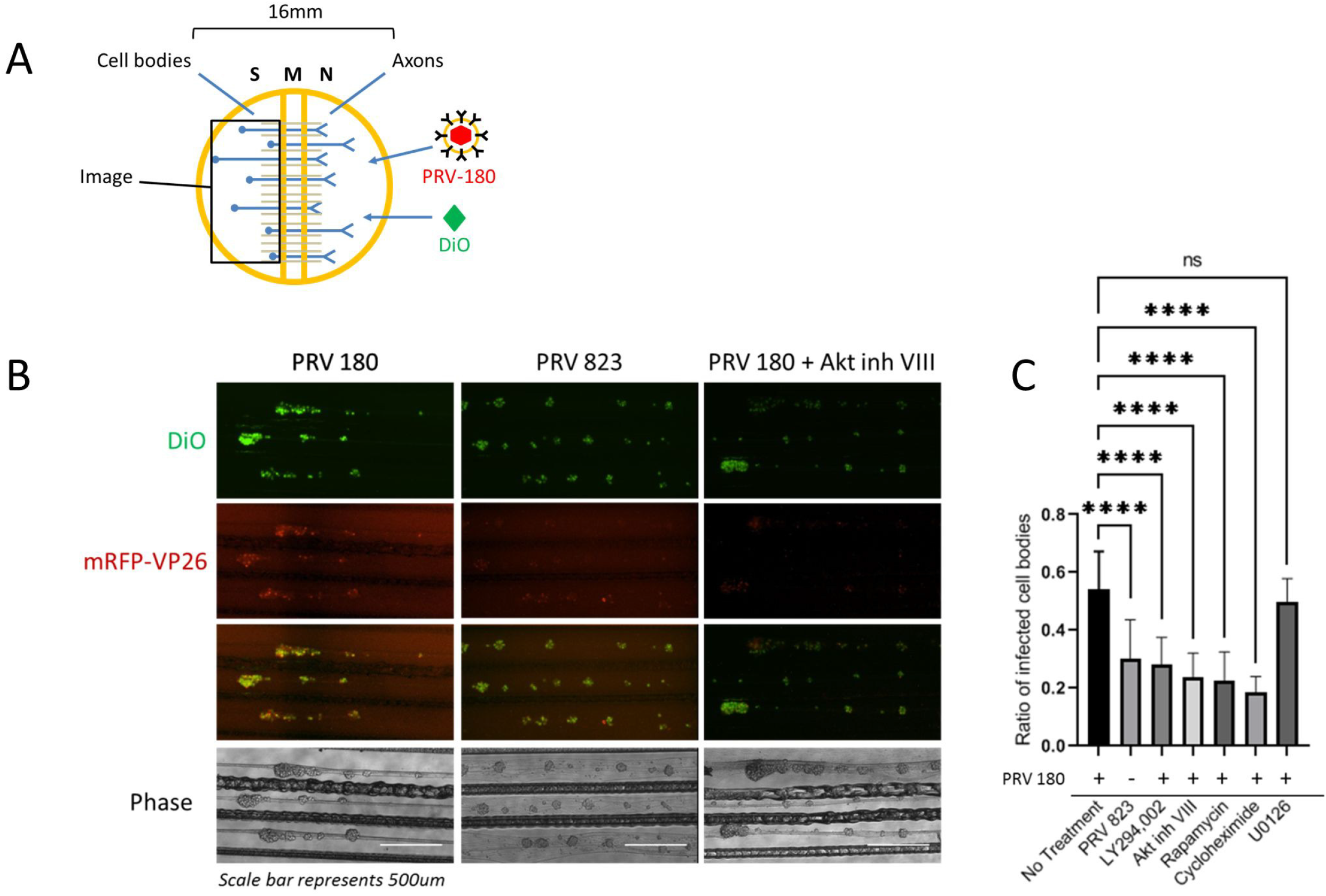
Disruption of Akt signaling pathways reduced PRV retrograde infection. (A) N compartment axons were treated with FAST-DiO 12hr prior to infection. PRV 180 or PRV 823 were added to axons and at 12hpi S compartment cell bodes were tile-imaged. For conditions involving inhibitor treatment, inhibitor was added to N compartments 1hr prior to infection and at 6hpi unabsorbed virus inoculum and inhibitor were removed from the N compartment. (B) Tile-images of neuron cell bodies in S compartments. Scale bar represents 500um. (C) Quantification of primarily infected cells. The ratio of dual-colored to total green cell bodies for each condition was calculated. Mean ± Std Dev with n = 10 chambers for each condition are plotted with *** p ≤ 0.001, using a one-way ANOVA (ns = not significant).

Local translation of axonal mRNA after PRV infection promoted the efficient retrograde transport of nucleocapsids through the axon ^10^. To determine if Akt-mToRC1 is also required for efficient transport, N compartment axons were infected with PRV 180 and at 2hpi, videos of fluorescent nucleocapsids trafficking through the M (middle) compartment were recorded (Fig.3A). Maximum intensity projections and kymographs were created from the videos of moving nucleocapsids to analyze transport kinetics (Fig.3B). Moving nucleocapsids were represented as continuous “tracks”. When N compartments were pretreated with LY294,002, Akt inhibitor VIII, rapamycin, or cycloheximide the number of moving nucleocapsids was reduced by ∼75.8% ± 24.4%, ∼76.3% ± 24.4%, ∼79% ± 22.8%, ∼75.8% ± 40.8% respectively compared to PRV 180 infection alone (Fig.3C). The net displacement of moving nucleocapsids (length of tracks) was also reduced by ∼50% when these inhibitors were present (Fig.3D). Nevertheless, the velocity of nucleocapsids that were moving in each condition was the same (Fig.3E), suggesting that engagement with the transport machinery, not the transport speed was disrupted. U0126 treatment had no significant effect on nucleocapsid transport or processivity. Taken together, these data demonstrate that Akt-mToRC1 signaling is induced by PRV infection in axons and is required for efficient retrograde transport of nucleocapsids. These effects were comparable to those observed when translation was blocked, suggesting Akt-mToRC1 activation acts early after PRV infection to promote translation in axons.

**Fig 3.**
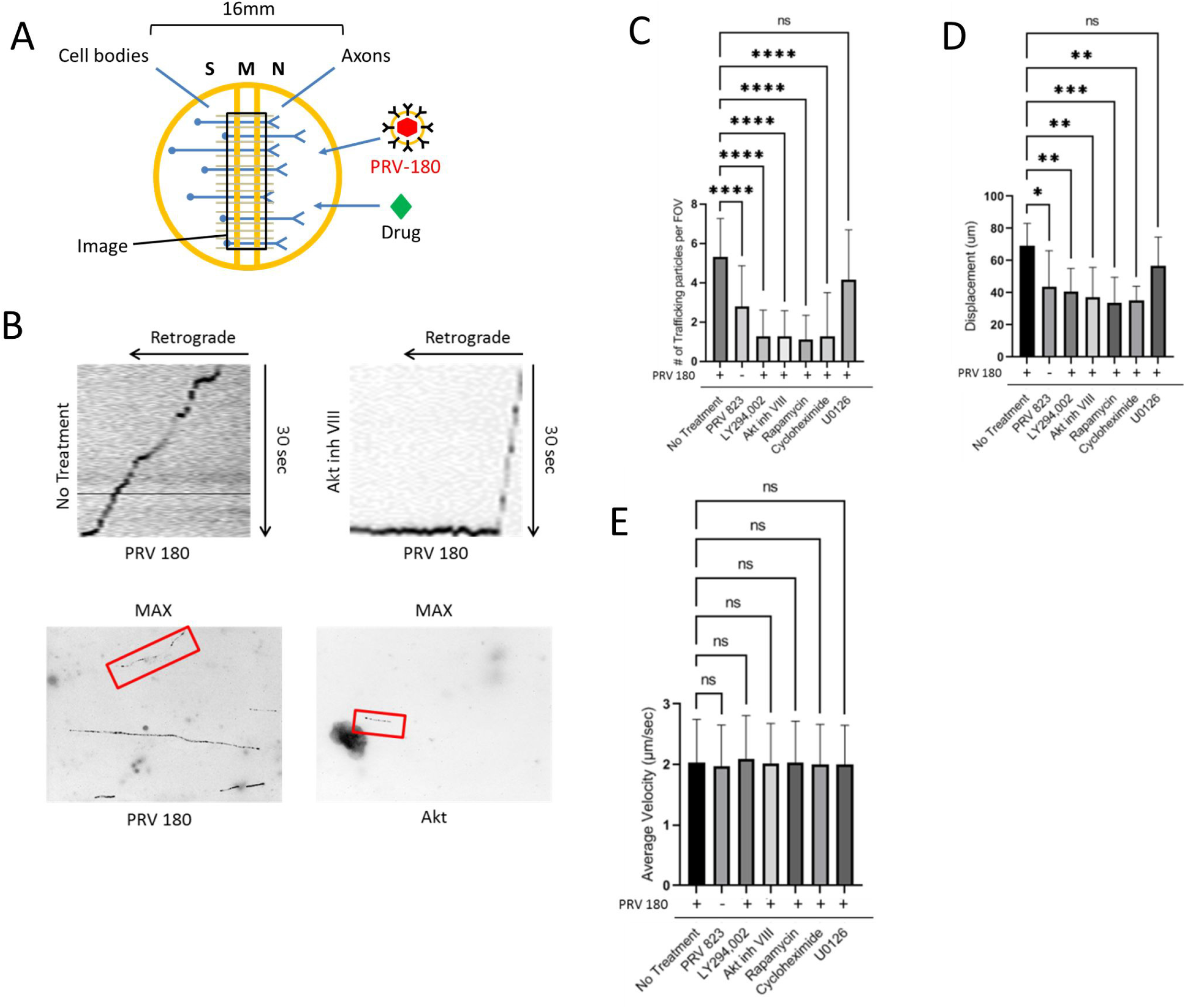
Quantification of PRV transport kinetics in axons. (A) N compartments were infected with PRV 180 or PRV 823. 2hpi 30sec videos of moving nucleocapsids in M compartments were recorded. Inhibitor was added to N compartments 1hr prior to infection when specified. (B) Maximum intensity projections (bottom) were created from the videos to visualize nucleocapsid displacement. Moving nucleocapsids are represented as tracks in the image (red box). Kymographs (top) were made from the maximum intensity projections to visualize nucleocapsid velocity throughout the recording process. Diagonal lines starting from the upper right corner represent retrograde movement. Horizontal lines represent stationary nucleocapsids. (C) Quantification of the number of moving nucleocapsids in M compartments. (D) The displacement of individual nucleocapsids over the 30sec recordings were measured for each condition. (E) The average velocity of the nucleocapsids moving in the retrograde direction were calculated by acquiring the mean of all instantaneous velocities ≥ 1um/sec. Data represent mean ± Std Dev with n = 7 chambers and 5 fields of view per chamber. **** p ≤ 0.0001, *** p ≤ 0.001, ** p ≤ 0.01, * p ≤0.05 using a one-way ANOVA (ns= not significant).

### Akt phosphorylation in axons requires PRV US3 ser/thr kinase

HSV-2 induces Akt phosphorylation upon binding of the glycoprotein gB on the virion envelope with α_v_β_3_ integrins leading to release of intracellular calcium stores to promote entry of nucleocapsids into the cytoplasm ^13^. We determined if virion binding to the cell surface or fusion of the viral and cellular membranes were responsible for Akt phosphorylation and if Akt phosphorylation affected transport directly or indirectly by regulating entry of nucleocapsids into the cytoplasm.

N compartment axons were infected with PRV-Becker mutants lacking either glycoprotein D (gD) (PRV GS442), the viral envelope protein required for specific PRV binding to the nectin-1 cell receptor, or glycoprotein B (gB) (PRV 233), the viral envelope protein required for fusion of the viral and cell membranes. Akt phosphorylation was monitored via western blot (Fig.4) and was unchanged after infection with either of these mutants suggesting that both binding and membrane fusion are required for Akt phosphorylation.

**Fig 4.**
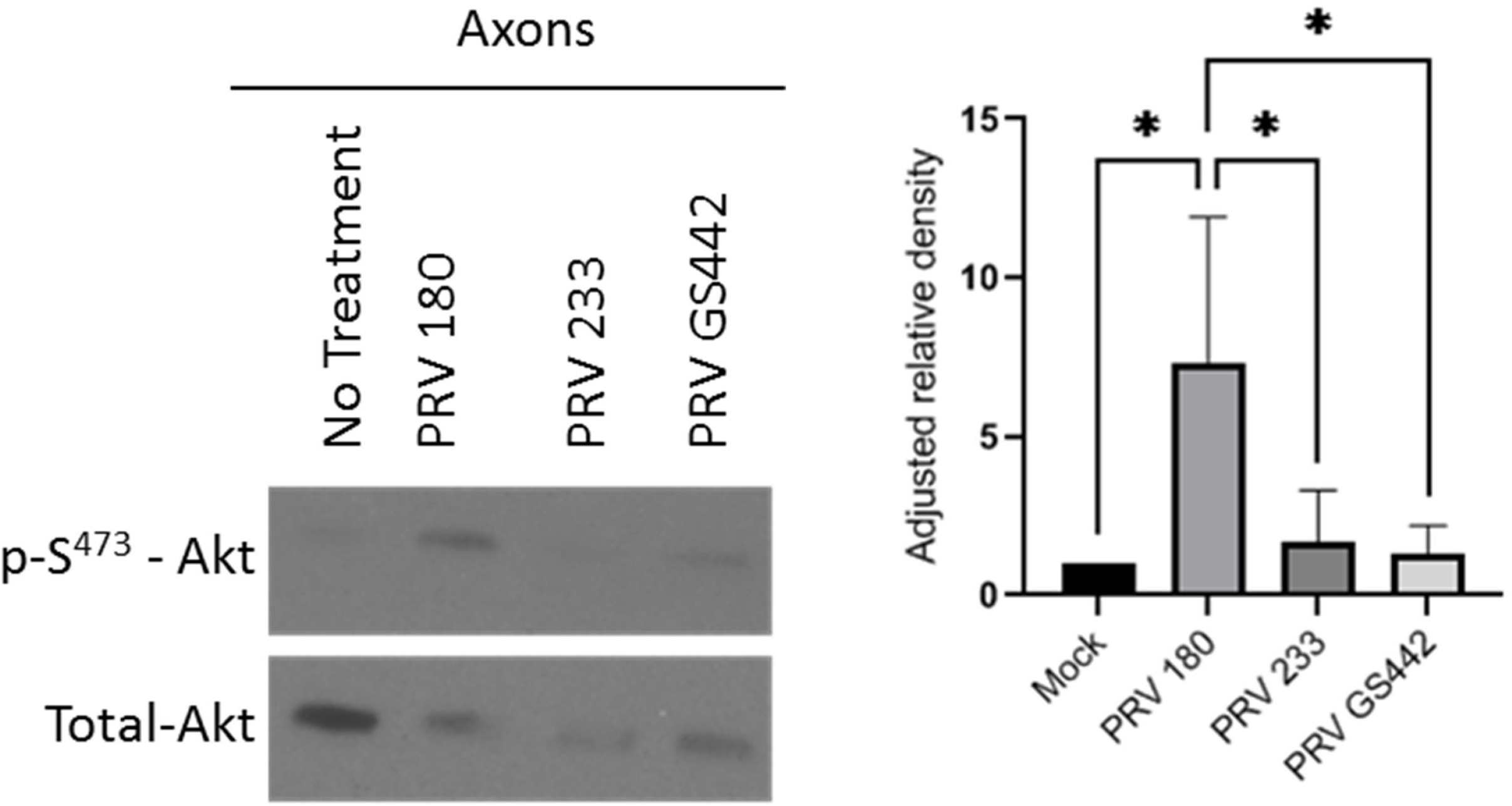
Akt phosphorylation in axons occurred after entry of PRV into the cytoplasm. Immunoblot of p-S^473^-Akt in axons infected with PRV 180, PRV 233, or PRV GS442 in N compartments. p-S^473^-Akt bands were normalized to total-Akt bands using a densitometry assay. Mean ± Std Dev with n = 3 for each condition are plotted with * p ≤ 0.05, using a one-way ANOVA.

To determine if Akt phosphorylation was induced by PRV after entry into the cell, we investigated the role of US3. US3 is an AHV inner-tegument protein with ser/thr kinase function^19^. US3 induces phosphorylation of Akt (in PRV) and downstream Akt substrates (in HSV-1) ^14,18^. When we infected axons with a PRV-Becker mutant lacking US3 (PRV 813NS) or a mutant lacking a functional kinase domain (PRV 815KD), no Akt phosphorylation was observed (Fig.5). When axons were infected with a revertant of PRV 813NS, termed PRV 813R, Akt phosphorylation was restored (Fig.5). These data indicate that US3 is responsible for inducing Akt phosphorylation after PRV infection in axons.

**Fig 5.**
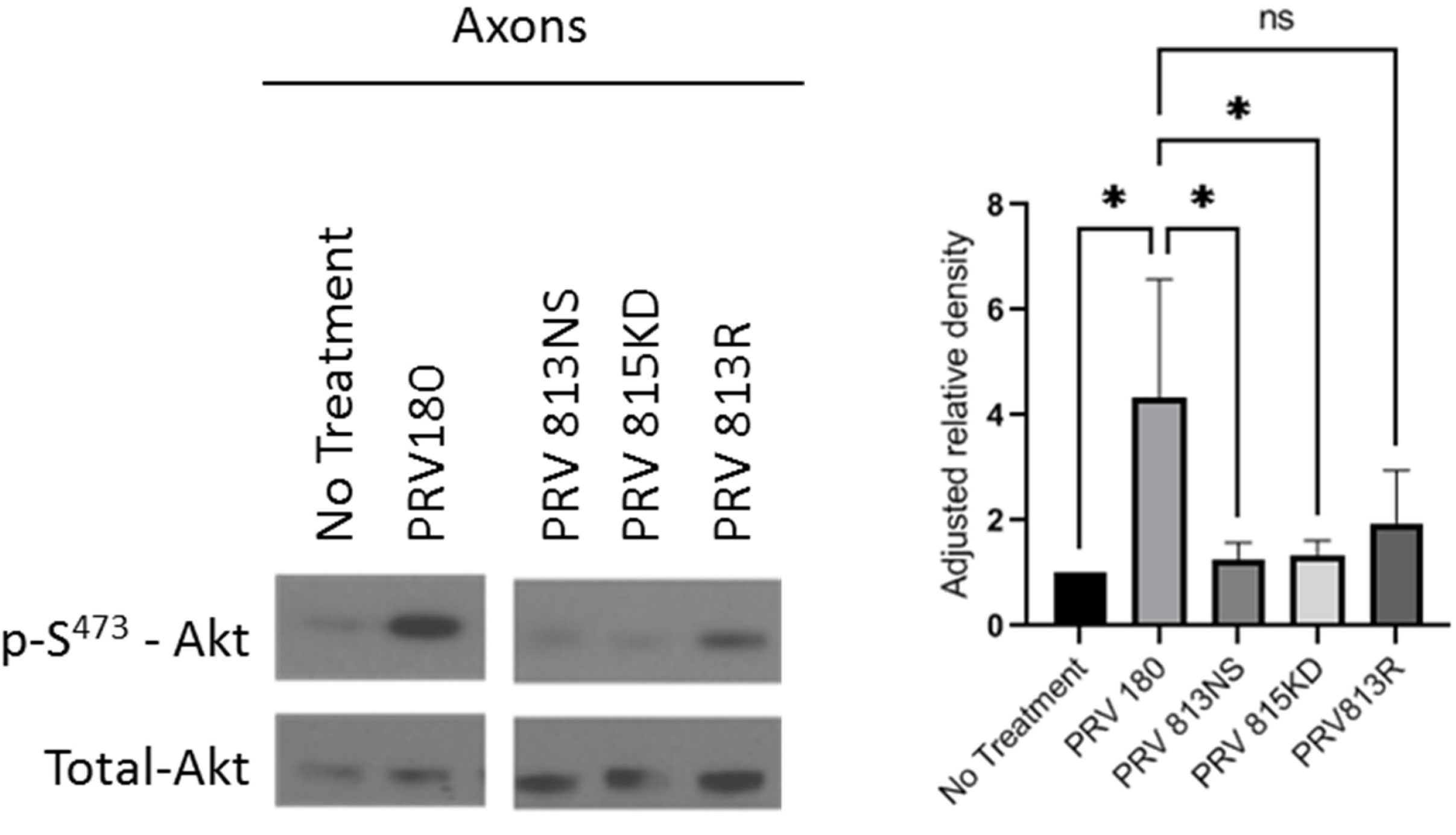
US3 induced Akt phosphorylation in axons. Immunoblot of p-S^473^-Akt in axons infected with PRV 180, PRV 813NS, PRV 815KD or PRV 813R in N compartments. p-S^473^-Akt bands were normalized to total-Akt bands using a densitometry assay. Mean ± Std Dev with n = 3 for each condition are plotted with * p ≤ 0.05, using a one-way ANOVA.

To determine if US3, and by extension Akt phosphorylation, are required for entry of PRV into cells, rat fibroblasts (Rat2) were infected with multiplicity of infection (MOI) of 5 of either PRV 180 or a ΔUS3 mutant with the mRFP-VP26 fusion (PRV823) at 4C, a temperature permissive to binding, but not entry. 1hpi cultures were washed to remove inoculum and the temperature was brought up to 37°C to allow for entry. After 15 minutes cells were washed with a low pH citrate buffer (pH 3) to inactivate any un-entered virus particles and the infection was permitted to continue for another hour. Cells were fixed and nucleocapsid fluorescence was visualized by fluorescence microscopy (Fig.6). Nucleocapsids present within the cytoplasm were counted manually and no significant difference was seen indicating US3 and Akt phosphorylation do not affect PRV entry in these cells.

**Fig 6.**
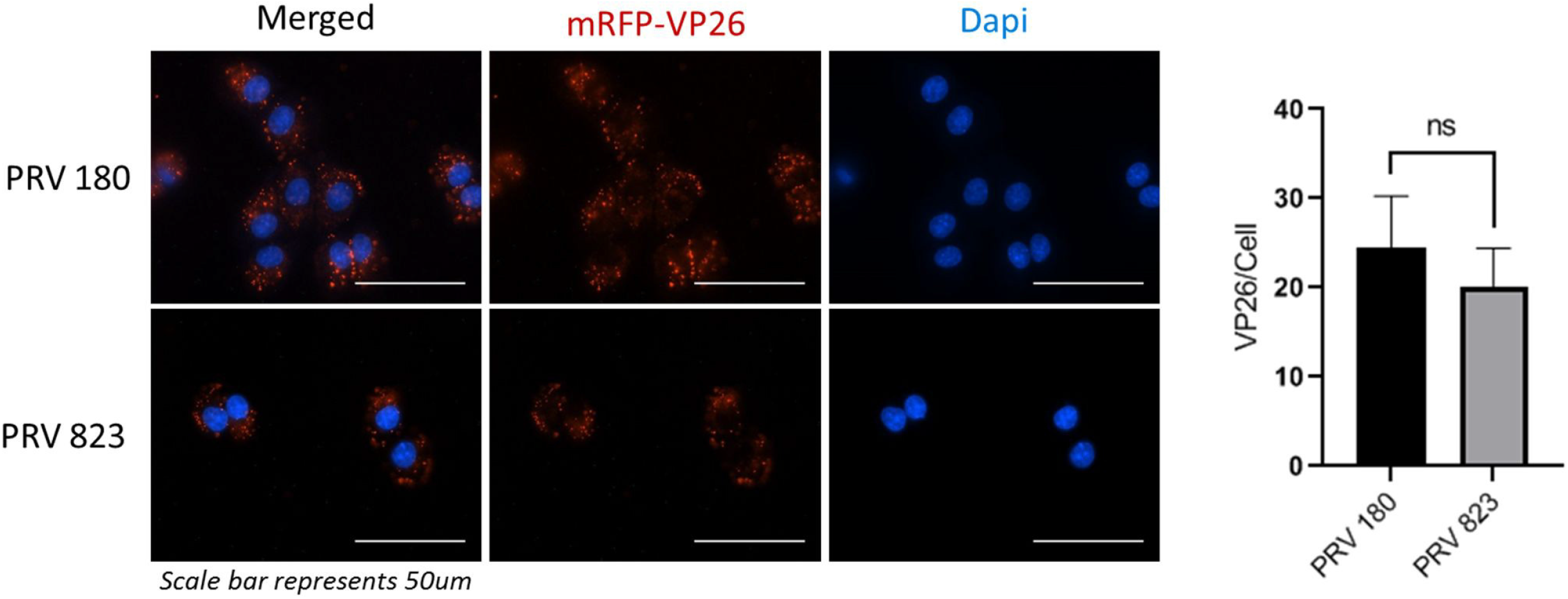
US3 does not affect entry of PRV into cells. Fluorescence imaging of PRV 180 and PRV 823 nucleocapsids in Rat2 fibroblasts. A synchronized infection assay was used to allow entry of all bound virion particles to occur simultaneously. Nucleocapsids that entered the cytoplasm were manually counted. Data represents mean ± Std Dev with 5 replicates and 2-5 cells per replicate for each condition using an unpaired t-test (ns = not significant).

Next, we determined if US3 is required for efficient retrograde infection of neurons using the technique described in Fig.2A. Compared to infection with PRV 180, infection with PRV 823 in axons led to a ∼44.7% ± 23.8% reduction in dual-colored cell bodies in the S compartment (Fig.2B). PRV 823 infection in axons in the N compartment also led to a ∼47.3% ± 38.4% reduction in the number of trafficking nucleocapsids in the M compartment when compared to PRV 180 (Fig.3C). PRV 823 transport kinetics were comparable to those of PRV 180 when Akt-mToRC1 signaling or translation was disrupted (Fig.3D, E,); net displacement was significantly reduced (∼37% ± 31.2%), but not transport velocity. Taken together, these data suggest US3 mediates efficient retrograde transport of PRV nucleocapsids through axons by inducing an Akt-mToRC1 signaling pathway.

### US3 and Akt phosphorylation are required for virus-induced local translation in axons

We directly tested whether US3 and Akt phosphorylation were required for translation by performing a surface sensing of translation (SUnSET) assay. Axons in the N compartment were either untreated or treated with Akt inhibitor VIII, cycloheximide, or U0126 then infected with PRV 180 or PRV 813NS. Puromycin was added to both N and S compartments to label nascent peptides 15 minutes prior to harvesting for western blot (Fig. 7A). Nascent peptides were visualized using a monoclonal puromycin antibody. Infection of axons with PRV 180 but not PRV 813NS led to an increase in puromycin incorporation when compared to mock, indicating that US3 is required for PRV-induced translation in axons. When Akt phosphorylation and translation were inhibited, puromycin incorporation did not increase past mock level indicating Akt phosphorylation is also required for translation to occur. Pretreatment with U0126 had no effect on puromycin incorporation showing the Ras/MAPK pathway does not affect PRV-induced translation in axons. (Fig.7B).

**Fig 7.**
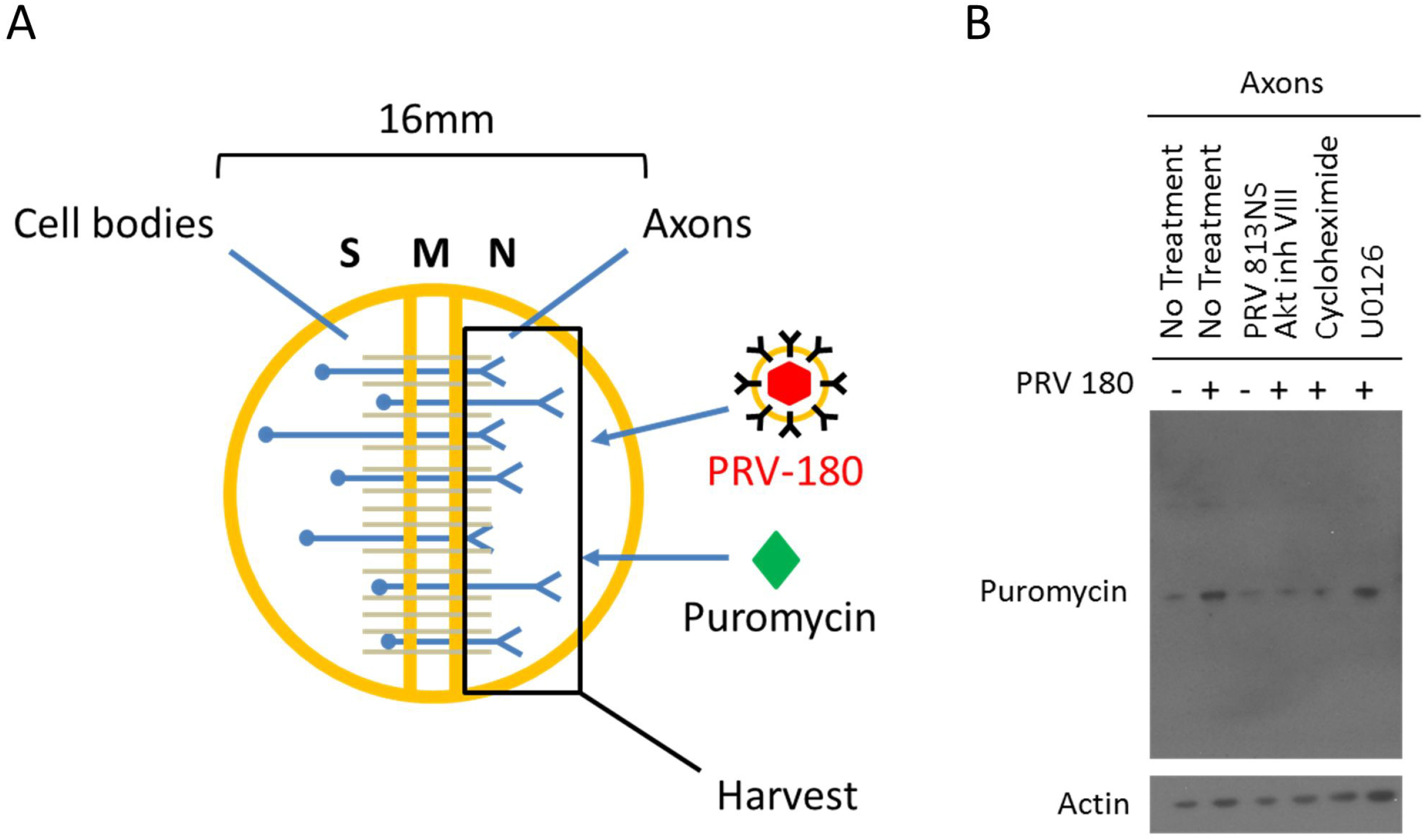
US3 and Akt phosphorylation are required for virus-induced translation in axons. A) N compartments were infected with PRV 180 for 1hr prior to harvest. Puromycin was added to N compartments at 45mpi to label nascent peptides. B) Immunoblot of puromycin incorporated peptides from axons in the N compartment. PRV 180 or PRV 813NS were added to N compartments for 1hr. Puromycin was added to N compartments 45mpi. Inhibitor was added 1hr prior to infection when specified. A single band is visible for each condition in this blot, representing the most abundant peptide synthesized at the time of incubation. Other peptide bands become visible at higher exposure times.

## Discussion

PNS neurons are highly polarized; with axon terminals extending centimeters away from their cell bodies. Accordingly, the transport of molecular messengers over these long distances is highly regulated ^37^. Axons contain within them the complete set of protein synthesis machinery, including multiple subsets of localized mRNA that are used to synthesize proteins in response to changes in the extracellular environment and send messages to the cell body to produce a global response to stimulus ^38,39^. PRV hijacks these processes to promote its efficient retrograde transport from the site of infection, in the axon, to the cell body ^10^. In this study, we have discovered that PRV utilizes an Akt-mToRC1 signaling pathway to induce translation in axons at early time points after infection. The inner-tegument protein kinase, US3, is required for the induction of this signaling pathway and acts upstream of PI3K to promote Akt phosphorylation (Fig.8).

**Fig 8.**
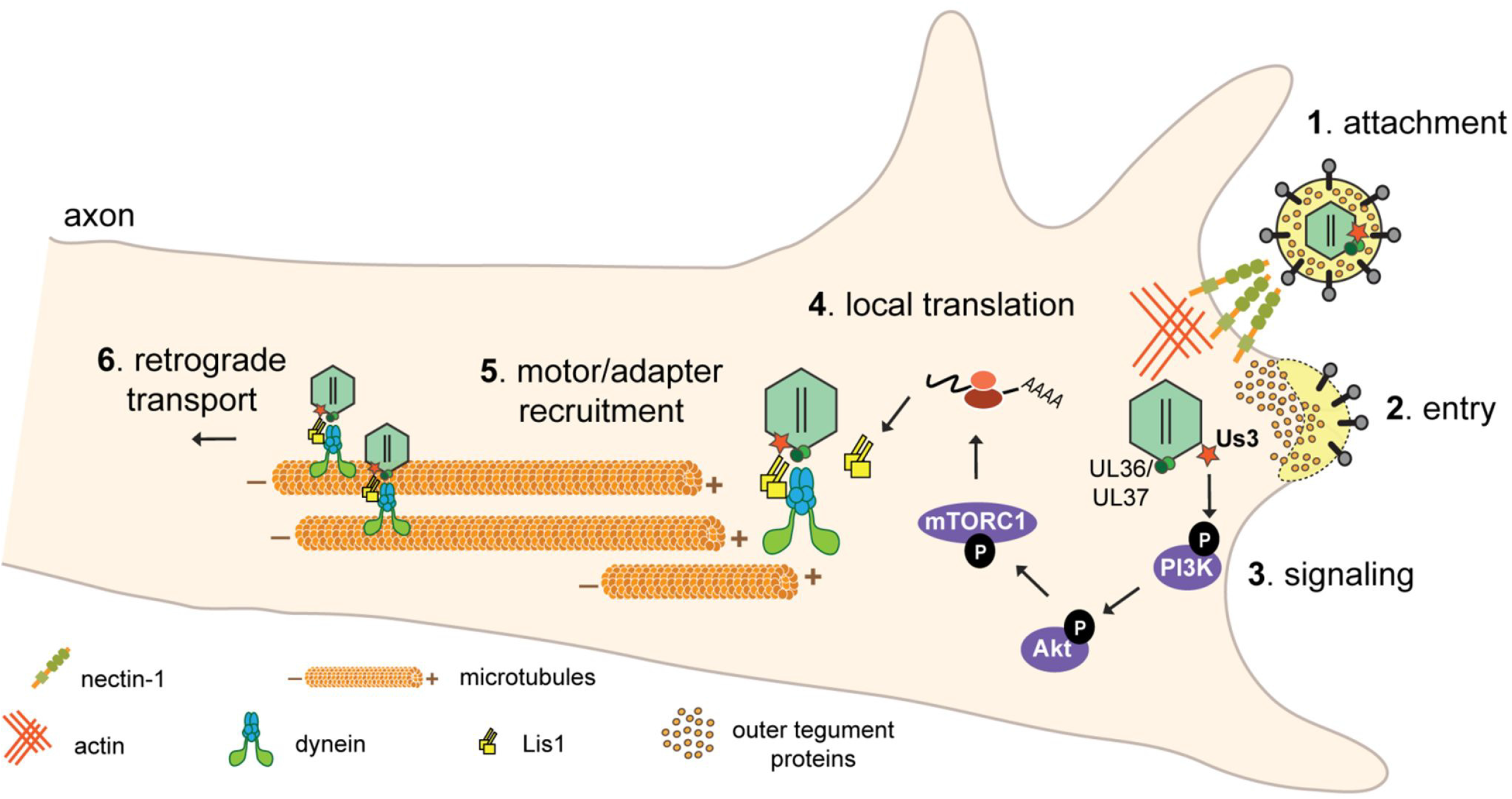
The model PRV-induced translation in axons. 1) PRV virions bind to nectin-1 receptors on the host-cell membrane. 2) Entry is mediated by fusion of the viral envelope with the cell’s plasma membrane allowing nucleocapsid and tegument proteins to enter the cytoplasm. Inner tegument proteins (US3, UL36, and UL37) stay bound to nucleocapsids. 3) US3 stimulates a PI3K/Akt-mToRC1 signaling pathway to induce 4) translation of axonal mRNAs leading to Lis1 expression. 5) Nucleocapsid engagement with the retrograde transport machinery is mediated by the inner tegument and Lis1 and 6) subsequent retrograde transport through the axon occurs.

Infection of axons with PRV 180 in the presence of the PI3K inhibitor, LY294,002, significantly reduced the number of moving nucleocapsids (Fig. 3C), and reduced the distance nucleocapsids traveled before stopping (Fig.3D). US3 must act upstream of PI3K to promote efficient transport, but it is unknown if PI3K is the direct kinase target of US3. Earlier studies have shown that US3 is responsible for the reorganization of the actin cytoskeleton via cofilin activation ^6^. US3 interacts directly with group 1 PAKs (p-21 activated kinases) to promote cofilin dephosphorylation (activation), leading to disassembly of filamentous actin and the entry of virion tegument and nucleocapsid into the cytoplasm ^40^. Group 1 PAKs play an important role in cytoskeleton rearrangement and apoptosis signal transduction ^41–48^ and have been shown to interact with various members of PI3K and Akt signaling pathways ^49^. It is possible that the cross-talk between group1 PAKs and the PI3K-Akt signaling pathway could be utilized by US3 to promote cytoskeletal rearrangements and local translation.

When US3 was absent or PI3K/Akt-mToRC1signaling was disrupted, we observed a significant reduction in the number of transporting nucleocapsids and their processivity (Fig.3). Under these conditions, nucleocapsids moved shorter distances before stopping; however the velocity of these nucleocapsids while in motion was similar to wild-type infection without inhibitor present. This suggests nucleocapsids had difficulty engaging with the transport machinery rather than difficulty transporting once engaged. These results were similar to what was seen for PRV and HSV-1 mutants when the R2 domain of UL37 inner-tegument protein was altered ^4^. The UL37-R2 mutant replicates well in peripheral tissue but is unable to move efficiently in peripheral neurons. As a result, the mutant cannot establish life-long infection in the host, a property that would provide a new possibility for vaccine design ^4^. It’s possible that the use of a US3-null-UL37-R2 mutant would provide a similar or additive effect by further reducing the chance of retrograde infection in peripheral neurons.

Long-distance transport in axons requires the microtubule network and associated motor proteins. The kinesin motor proteins mediate anterograde (plus-end directed) transport and dynein motor proteins mediate retrograde (minus-end directed) transport ^50^. Dynein is a multisubunit complex composed of two heavy-chains, two intermediate chains, and two light chains that regulate motor activity and interactions with cellular cargo ^51–54^. To achieve spatiotemporal regulation of transport, various accessory factors associate with the dynein complex; one such factor is dynactin^50^. The UL36 inner-tegument protein interacts with dynactin to recruit nucleocapsids to dynein^8,9^; however this interaction alone is not sufficient to promote efficient transport. Dynein must be phosphorylated to obtain an active conformation^50,51,55–57^. Recently, it was shown that upon infection of epithelial cells with HSV-1, dynein intermediate chain 1B was phosphorylated at position S80 in an Akt and PKC independent manner ^58^. We have also observed dynein phosphorylation (at position T88) during PRV infection that was US3 dependent (A. D. Esteves and L. W. Enquist, unpublished data). US3’s kinase function could be mediating the activation of dynein throughout the retrograde transport process. More work needs to be done to determine the cause and function of dynein phosphorylation during AHV infection.

Axonal infection with PRV mutants lacking either gD or gB, the glycoproteins that mediate virion binding to the plasma membrane and carry out membrane fusion, were unable to stimulate Akt phosphorylation (Fig.4). These observations suggested that Akt is phosphorylated after entry of the nucleocapsid and tegument proteins in the cytoplasm. These findings are different from what was observed after HSV-1 and HSV-2 infection. Infection with these AHVs lead to the stimulation of a low-level Ca^2+^ fluctuation following binding of gD or gB to the nectin-1 co-receptor and heparin sulfate proteoglycans on the cell surface. This is followed by an interaction between Akt and gB leading to Akt phosphorylation, a larger Ca^2+^ fluctuation and entry of the nucleocapsid and tegument into the cytoplasm ^13^. We have shown that PRV nucleocapsid entry does not depend on US3, and by extension, Akt phosphorylation (Fig.6). However, we have not investigated the potential for the induction of Ca^2+^ fluctuation during infection and whether or not it participates in the retrograde transport process. Future work needs to be done to determine the effects of Ca^2+^ signaling in PRV infection.

Our work has revealed a novel role for US3 in the promotion of efficient microtubule-based transport of PRV nucleocapsids in axons. We determined that US3 signals through an Akt-mToRC1 signaling pathway, upstream of PI3K, to induce translation of localized, repressed mRNAs, a process required for efficient retrograde transport. The absence of US3 or the disruption of Akt-mToRC1 not only reduced the number of transporting nucleocapsids but also negatively impacted the processivity of nucleocapsids moving through the axon, suggesting engagement with the dynein-motor complex was disrupted. This work helps to clarify the viral and cellular factors involved in PRV entry and retrograde transport in axons and identifies US3 as a potential target for therapies aiming to prevent the spread of AHVs in the nervous system.

## Acknowledgements

This work was supported by the US National Institute of Health grant (5R37NS033506) to Lynn W. Enquist. The authors declare that they have no competing interests.

We thank Kevin Pfister for providing the phospho-dynein antibodies used in this study and Heath E. Johnson for assistance with particle tracking automation.

## Materials and Methods

### Primary neuronal culture

Superior cervical ganglia (SCG) neurons were isolated from embryonic day 17 Sprague-Dawley rat embryos and cultured as previously described ^29^. Briefly, SCG were trypsinized and mechanically dissociated. 35mm cell culture dishes were coated with poly-DL-ornithine (Sigma-Aldrich) and laminin (Invitrogen) then fitted with a Campenot tri-chamber (CAMP320 isolator rings, Tyler Research) using autoclaved silicone vacuum grease as an adherent. Dissociated SCG were seeded in one compartment (the S-compartment) of the tri-chamber filled with neurobasal medium (Life Technologies; 21103049) + 50x B-27 supplement (Life Technologies; 17504044) + 100x Penicillin-Streptomycin-Glutamine (ThermoFisher; 10378016) + 80ng/ml NGF (ThermoFisher; 13257019) and left to grow for ∼3weeks with media changes every 7 days.

### Cell lines and virus stocks

Porcine kidney epithelial cells (PK15, ATCC), and Rat2 cells (ATCC) were maintained in Dulbecco modified Eagle medium (DMEM, Hyclone) + 10% fetal bovine serum (FBS, Hyclone) + 1% penicillin-streptomycin (Hyclone). Virus propagation and titer were performed in PK15 cells unless otherwise specified. Rat2 cells were used for synchronized infection assays. LP cells (gB complementing cells derived from PK15 cells) were used to grow PRV 233 virus stocks (Lisa Pomeranz, personal communication). 100ug/ml Geneticin (G418) (InVivo Gen, ant-gn-1) was added to culture every 5^th^ passage to maintain gB expression. G5 cells (gD complementing cells derived from PK15 cells) were used to grow PRV GS442 virus stocks ^59^. 2.5mM L-histidinol dihydrochloride (Sigma, H6647) was added to cultures every 5^th^ passage to maintain gD expression. All cells were incubated at 37°C, and 5% CO_2_.

Unless otherwise specified, virus infections of PK15 and Rat2 cells were performed with DMEM + 2% FBS. Titers of virus stocks were determined as Plaque forming units (PFU). For live-cell imaging PRV 180 and PRV 823 stocks were used within 2 weeks of production for no more than one freeze-thaw cycle to preserve the fluorophore intensity. Virus stocks used are as follows:

| Virus Strain | Genotype | Reference |
| --- | --- | --- |
| PRV 180 | Becker wild-type strain with mRFP-VP26 fusion | 60 |
| PRV 813NS | US3-null Becker strain; nonsense mutation | 20 |
| PRV 813R | Revertant of 813NS | 20 |
| PRV 823 | US3-null Becker strain, mRFP-VP26 fusion | 20 |
| PRV GS442 | gD-null Becker strain, diffusible GFP expression | 59 |
| PRV 233 | gB-null Becker strain, diffusible GFP expression | 61 |

### Antibodies and chemicals

All antibodies were diluted in TBS-T (Tris buffered saline-Tween, 0.1%) treated with 3% bovine serum albumin (BSA) and stored in −20°C unless otherwise specified. Rabbit phospho-Akt (ser473)(D9E) (Cell Signaling Technology, 4060) was used at 1:1,000 for western blot (WB). Rabbit Akt antibody (Cell Signaling Technology, 9272) was used at 1:1,000 to detect total Akt in WB. Monoclonal anti-beta-actin antibody (Sigma Aldrich, A1978) was used at 1:10,000 for WB. Mouse monoclonal US3 7H10.21 ^20^ was used at 1:1,000 for WB. Rabbit phospho-dynein-threonine88 was a gift from Kevin Pfister and was used at 1:300 for WB. Mouse recombinant anti-cytoplasmic dynein intermediate chain (74.1) (Abcam, ab23905) was used at 1:500 for WB. Mouse monoclonal anti-puromycin clone 4G11 (EMD Millipore, MABE342) was used at 1:2,000 for WB.

All drug treatments occurred 1hr prior to infection, concentrations represent final experimental concentrations, and drugs were resuspended in DMSO unless otherwise specified. LY294,002 (Cell Signaling Technology, 9901) was used at a concentration of 20uM. Akt inhibitor VIII (Sigma, 124018) was used at a concentration of 5uM. Rapamycin (Sigma, 553210) was used at a concentration of 100uM. U0126 (Sigma, 662009) was used at a concentration of 10uM. Puromycin (AG Scientific, P-1033-SOL) was solubilized in deionized H_2_O and used at a concentration of 1ug/ml. FAST-DiO (ThermoFisher, D3898) was used a concentration of 5ug/ml and incubated for 12hr prior to infection (24hr prior to imaging).

### Western Blotting

SCG neurons in tri-chambers were seeded at a density of 1 SCG per S compartment. Dishes were washed 3x with warm PBS (phosphor buffered saline) then incubated with neurobasal media + 100x penicillin-streptomycin-glutamine (note the lack of B-27 supplement) for 12hr prior to infection. Either axons in the N compartment or cell bodies in the S compartment were lysed in the chamber with 40ul of 2x Laemmli buffer prepared from a dilution of 5x Laemmli buffer (10% SDS, 300mM Tris-Cl pH = 6.8, 0.05% bromophenol blue,100mM DTT, and 50% Glycerol in ddH_2_O). Lysates were boiled at 90°C for 5min then cooled on ice. Proteins were separated by SDS-PAGE and 4-12% gradient NuPAGE Bis/Tris gels. Proteins were transferred to nitrocellulose membranes (GE Healthcare, 45-0040002) using a Trans-Blot SD semi-dry transfer cell (Bio Rad). Membranes were blocked using 1x Pierce Clear Milk Blocking Buffer (ThermoFisher, 37587) for 30min at room temperature (RT) then washed 3x with TBS-T. Membranes were incubated in primary antibody dilution over night at 4°C followed by 3x TBS-T washes then incubation with horseradish peroxidase-conjugated secondary antibody (1:10,000 dilutions in 3% BSA-TBS-T) for 45min at RT and a final 3x washes with TBS-T. Chemiluminescent substrate, Supersignal West Pico (ThermoFisher, 34080), West Dura Extended Duration Substrate (ThermoFisher, 34075), West Femto Maximum Sensitivity Substrate (ThermoFisher, 34094), or West Atto Ultimate Sensitivity Substrate (ThermoFisher, A38554) were added to the membranes for 5min at RT. Protein bands were visualized by exposing the membranes on HyBlot CL autoradiography film (Denville scientific, E3018).

### SUnSET assay for labeling nascent peptides

Puromycin was added to the chamber compartment to be analyzed 15 minutes prior to harvest. The rest of the protocol follows the same as for WB. Puromycin-labeled peptides were detected with puromycin antibodies. See “Antibodies and chemicals” for the primary antibody used.

### Synchronized infection assay

Rat2 cells in culture were cooled to 4°C. PRV 180 or PRV823 inoculumns were added at an MOI of 5 and virion absorption to cells was allowed to occur for 1hr at 4°C before media was aspirated and cells were washed with chilled PBS 3x. Chilled 2% FBS DMEM was added to the dish and temperature was brought up to 37°C for 15 minutes to enable virion entry. Media was removed and infected cells were washed once with a low pH citrate buffer for 2min to inactivate virions that had not entered cells. Citrate buffer was removed and cells were washed 3x with PBS at RT before warm 2% FBS DMEM was added. Infected cells were incubated for 1 hour at 37°C.

Cells were fixed using 4% paraformaldehyde (PFA) for 10min at RT and permeabilized with 0.1% Triton x-100 (Sigma, T8787) for 10min at RT. After permeabilization, plates were blocked for 1hr at RT with 1% BSA in PBS, washed 3x with PBS then treated with DAPI (ThermoFisher, 62248) to stain cell nuclei for 5min at RT in the dark with 1ug/ml DAPI in 0.1% BSA PBS. Cells were washed 3x with PBS then imaged. Total and average number of red-fluorescent nucleocapsids per cell in a single field of view (FOV) were manually counted using NIS Elements Advanced Research software (Nikon).

### Retrograde transport assay

SCGs were seeded at 2/3 of an SCG per S compartment. Fast-DiO was added to axons in the N compartment 12hr prior to infection. N compartments were either untreated or treated with LY294,002, Akt inhibitor VIII, rapamycin, cycloheximide, or U0126 1hr prior to infection. Axons in N compartments were infected with 10^6^ PFU of PRV. At 6hr post infection unabsorbed virus inoculum and drug in N compartment were washed out and replaced with fresh neurobasal medium. At 12hr post infection cell bodies in the S compartment were tile-imaged for mRFP-VP26 (red) and DiO (green). Total numbers of green and dual-colored particles (red and green) were manually counted (see synchronized infection assay). The ratio of dual-colored cell bodies to total green cell bodies was determined per S compartment and averaged for all conditions.

### Single particle tracking in tri-chambers

SCGs were seeded at 2/3 of an SCG per S compartment with optical plastic tissue culture dishes (Ibidi, 81156-400). N compartment axons were either untreated or treated with drug for 1hr prior to infection. Infections were initiated at 10^6^ PFU for 2hr before time-lapse imaging was used to visualize moving fluorescent virus particles in the M compartment. 30 second recordings of a single FOV were completed at approximately 2 frames per second. The total number of moving virus particles were counted. For instantaneous velocity measurements the TrackMate plugin for imageJ was used. Velocities were parsed into bins in a histogram ranging from - 4um/sec (the maximum anterograde velocity recorded) and +4um/sec (the maximum retrograde velocity recorded) with intervals of 1um/sec. Instantaneous velocities in bins 1um/sec to 4um/sec were averaged across all conditions to determine the average velocity of all moving virion particles. Velocities in the 0um/sec bin were assumed to be stationary particles and were not included in the average velocity measurements.

### Imaging processing and analysis

All imaging was conducted on a Nikon Eclipse Ti inverted epifluorescence microscope using a CoolSNAP ES2 CCD camera. Images and movies were processed using ImageJ ^62^ and NIS Elements Advanced Research software (Nikon). Comparative images were all captured with the same exposure times, brightness and contrast adjustments were applied to the entire image and alterations were applied equally across conditions.

To analyze trafficking virus particles in the M compartment, imageJ was used to create maximum intensity projections of all videos to visualize the path of moving particles. Particle trajectories were manually counted and taken to represent a single moving particle. The directionality and velocity of a moving particle was visualized using the imageJ multi kymograph tool. Particle lengths were calculated by measuring the total particle distance traveled over the course of the 30sec movie using the segmented line tool in imageJ. In order to be counted, particles must not enter or exit the FOV during the entire 30sec recording time.

### Statistical analysis

All data were analyzed using GraphPad Prism 9 (GaphPad Software, La Jolla California USA, www.graphpad.com). Figure legends provide the statistical test used for each analysis.

